# A vulnerability assessment framework for terrestrial invertebrate species threatened by wildlife trade

**DOI:** 10.64898/2026.11.11.766051

**Authors:** Charlotte R. Lassaline, Sean McGowan, Erinn Fagan-Jeffries, Adam Toomes, Phillip Cassey

## Abstract

Wildlife trade is an emerging threat to terrestrial invertebrates in Australia, yet its impact on native populations remains poorly quantified. Short range endemic (SRE) species, often characterised by restricted geographic ranges, low reproductive output, low vagility, and specialised habitat conditions, are particularly vulnerable to anthropogenic pressures, including over-exploitation via wild harvesting. Using a multi-year dataset of Australian invertebrate trade records, we identified species that are both utilised in the pet trade and exhibiting SRE traits, and assessed their risk of population decline by integrating trade frequency with species’ distribution, abundance, and biological vulnerability. We identified 266 species traded in Australia with an area of occupancy less than 500 km², with arachnids disproportionately represented among those at greatest relative risk of population decline from over-exploitation. While these rankings should be interpreted as a methodological demonstration rather than a definitive measure of extinction risk, they provide a transparent approach for screening species where trade pressure may intersect with biological vulnerability. To the best of our knowledge, this is the first tested approach to holistically assess this risk in invertebrates. We present a proof-of-concept framework for quantifying trade-related vulnerability in terrestrial invertebrates, providing a transferable approach that can be refined and applied across a wide range of taxa threatened by wildlife trade.

## Introduction

Human activities are altering the natural environment at historically unprecedented rates, driving widespread biodiversity loss and ecosystem change (Braby 2018; Branco and Cardoso 2020; Hald-Mortensen 2023). Predicting how species are affected by these pressures and how they will respond is essential for understanding broader consequences on ecosystem structure, function and resilience (Isbell et al. 2023). However, conservation action is often implemented reactively, only after species or populations exhibit clear signs of decline (Brooks et al. 2006; Cardoso et al. 2011). There is an urgent need to shift towards proactive conservation approaches that identify vulnerable species before population declines reach irreversible levels (Mamo et al. 2020; Simmonds et al. 2020).These approaches rely on predictive methods capable of assessing species vulnerability prior to documenting substantial ecological impact.

The need for vulnerability assessment methods is particularly acute for terrestrial invertebrates, which remain largely absent from conservation frameworks, despite facing numerous and escalating threats (Cardoso et al. 2011; dos Santos et al. 2020). Population declines in invertebrates may occur long before they are discovered or formally documented, limiting opportunities for conservation intervention. In Australia, invertebrates comprise approximately 98% of animal diversity and perform essential ecosystem functions, including pollination, decomposition, soil turn over, and pest control (Black et al. 2001; Cranston 2010; Eisenhauer et al. 2019; Wilson 1987). These processes underpin ecosystems of substantial ecological, economic, and cultural importance. However, Australia’s terrestrial invertebrates are increasingly threatened by multiple interacting pressures, including habitat loss, displacement by introduced species, pesticide use, climate change, and more recently, the growing demand for wild-caught specimens in pet trade (Figure 1) (Branco and Cardoso 2020; Lassaline et al. 2025; Woinarski et al. 2024). Despite their ecological importance and mounting threats, invertebrates remain critically underrepresented in scientific research and conservation policy, both globally and within Australia, where efforts remain disproportionately focused on vertebrate taxa (Cardoso et al. 2011; Clark and May 2002; Isbell et al. 2023; Lassaline et al. 2025).

**Figure 1:**
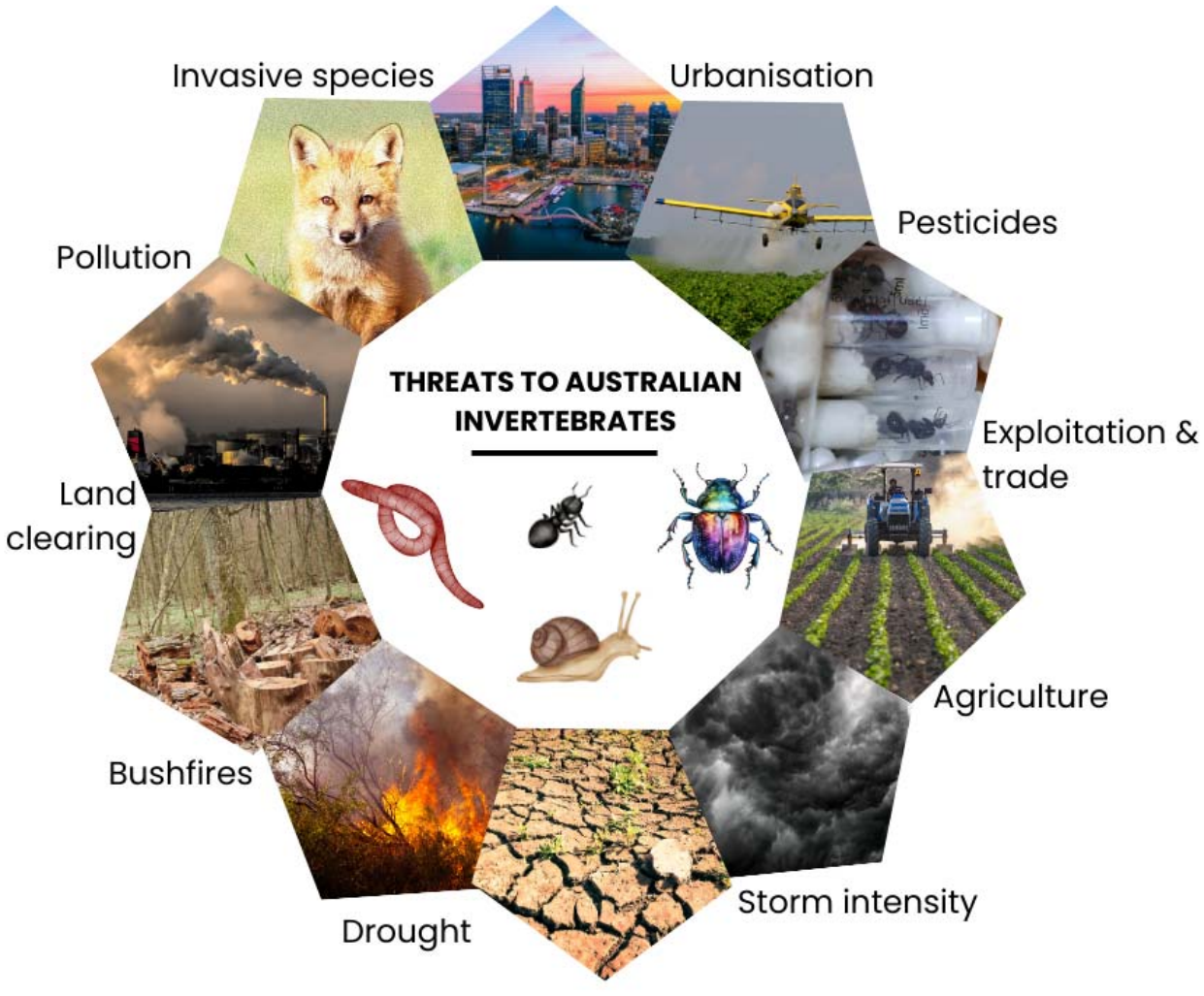
The extensive variety of threats faced by Australian terrestrial invertebrates (Isbell et al. 2023; Kehoe et al. 2021; Woinarski et al. 2024).

Australia is both a source and destination for traded invertebrate species, many of which are collected from the wild with little regulatory oversight (Lassaline et al. 2023; Lassaline et al. 2026a). Their cryptic behaviour, small body size, and immense taxonomic diversity, combined with Australia’s vast and sparsely human-populated landscape, make their exploitation particularly difficult to detect and monitor (Delu et al. 2026; Lassaline et al. 2026a). This challenge is compounded by taxonomic knowledge gaps, as many traded invertebrates remain undescribed and are more difficult to assess and recognise under formal conservation or legal frameworks which assume species-level names are available. Recent trade surveys within Australia have revealed a wide diversity of native terrestrial invertebrates sold as pets, with both the volume and the diversity of species increasing over time (Lassaline et al. 2026b). Consequently, highly vulnerable taxa may be exposed to exploitation through the pet trade before their conservation threats are even recognised. The Australian Commonwealth Environmental Protection and Biodiversity Conservation Act 1999 (EPBC Act), along with state and territory legislation, provides an important framework for species protection in Australia (Taylor et al. 2018). However, funding and resources are typically directed toward nationally listed species, which may leave regionally significant taxa with less support (New and Yen 2013). Conservation progress is further constrained by poor public visibility, and how funding is often only available for species already recognised as threatened (Cardoso et al. 2011; New and Yen 2013). As the Australian terrestrial invertebrate pet trade continues to grow, further research is needed to identify taxa most vulnerable to over-exploitation and inform conservation action, before populations are adversely impacted (Lassaline et al. 2026b).

Short-range endemic (SRE) species are organisms with limited geographical distributions, typically smaller than 10,000 km^2^ (Harvey 2002). They are often confined to localised or fragmented habitats, are poor dispersers with low vagility, rely on specialised environmental conditions, and have low reproductive outputs (Framenau and Harvey 2009). Short range endemism is widespread across terrestrial invertebrates, and in Australia several groups primarily consist of SREs, including Gastropoda (snails and slugs), Araneae (spiders, particularly mygalomorph spiders), and Diplopoda (millipedes) (Harvey 2002). These biological traits, when combined with anthropogenic threats such as exploitation through pet trade, place SREs at heightened risk of population decline and extinction.

Here, we present a framework for assessing the vulnerability of Australian terrestrial invertebrate species to overexploitation from domestic wildlife trade. Using previously collected trade data from Australian social media, pet stores, expos, and websites (Lassaline et al. 2026b), we identified species that may be at the greatest risk of population decline from exploitation pressure. We present a list of taxa potentially requiring urgent conservation attention and stronger consideration within wildlife trade regulation, and provide recommendations to better protect invertebrates impacted by exploitation. While our framework demonstrates an initial proof-of-concept to be guided by future refinement, validation, and broader assessment efforts, it provides a proactive approach for screening species that may face elevated trade-related risk before substantial population declines are documented. By supporting early recognition of vulnerable taxa, we seek to contribute towards more preventative conservation strategies for Australian fauna.

## Methods

Our species prioritisation assessment involved four key steps: 1) Scoping and planning, including procuring the data, identifying species to include in the assessment, and defining the sensitivity and exposure variables; 2) Assessment preparation, including calculating species distributions (area of occupancy; AOO), and collating information on their life-history traits and trade pressure; 3) Scoring variables by determining their relevant weighting using the analytic hierarchy process (AHP), involving pairwise comparisons undertaken by scientific experts; and 4) Calculating and analysing species vulnerability, presenting a conservation prioritisation list of the most vulnerable species (Figure 2).

**Figure 2:**
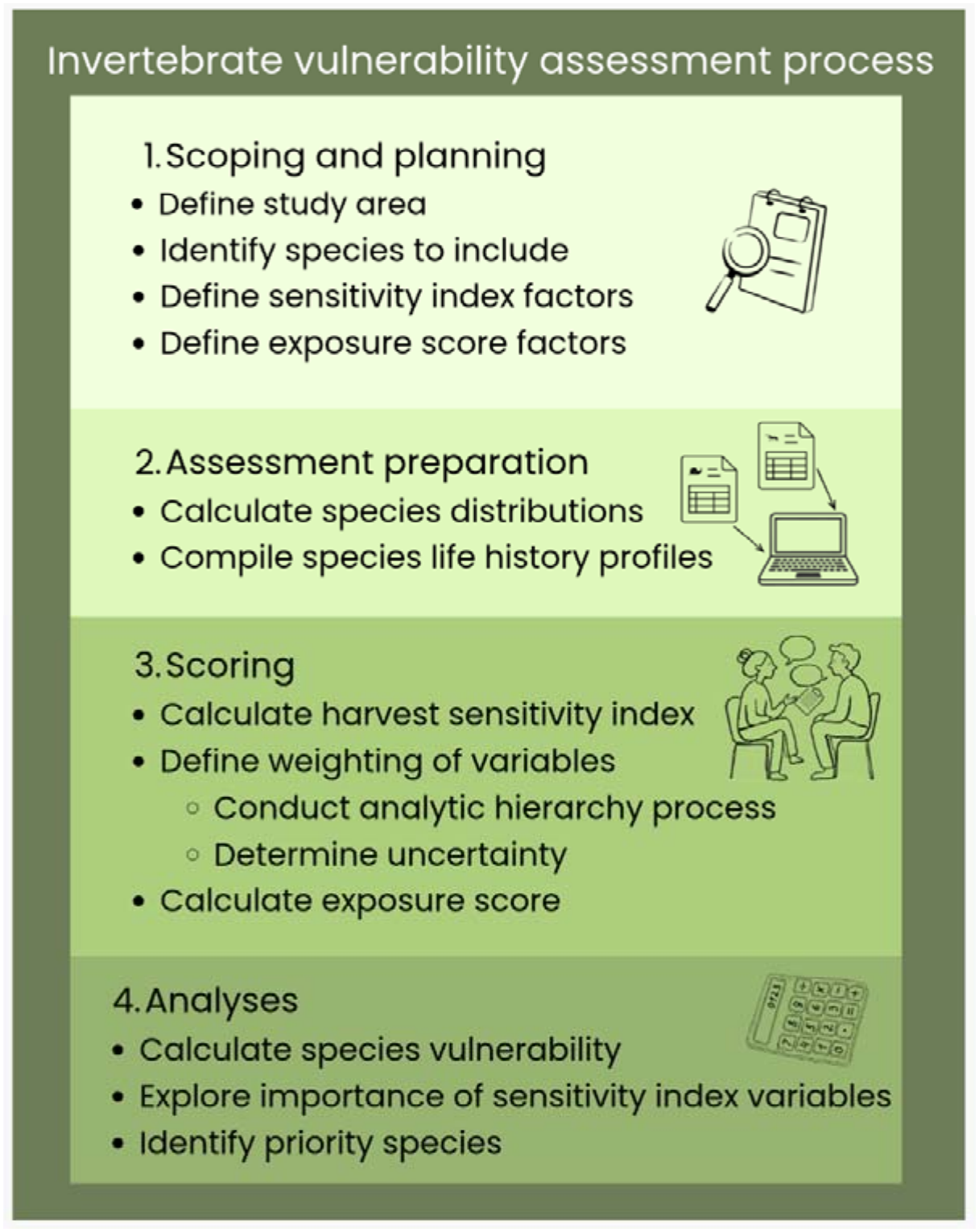
The four-step process for creating a species vulnerability assessment framework for terrestrial invertebrates. This broadly includes: (i) scoping the available data and organising the framework structure, (ii) collecting the available data, (iii) determining relative weighting for each variable through the analytic hierarchy process, and (iv) calculating the vulnerability index for each species.

### 1) Scoping and planning

Our considered species list considered was derived from research on the Australian terrestrial invertebrate trade, reporting 586 invertebrate species in domestic trade (Lassaline et al. 2026). This included species from trade surveys between 2019 and 2025, across Australian pet stores, online retailers, social media groups, public insect expositions, and a public classifieds’ website. We define the study area for this framework as the Australian mainland States & Territories as well as Tasmania.

While there are many sensitivity variables that influence a species vulnerability, and could be incorporated into future vulnerability assessments, our study was constrained by the substantial data deficiencies associated with terrestrial invertebrates. We did not include variables for which data was unavailable for more than 50% of species. Consequently, the selection of sensitivity variables was guided by both biologically relevant traits, and the availability of information. Our resulting framework ultimately proposes the Sensitivity index to be a combination of two categories: i) population and demography information; and ii) life history traits; proposed for similar vulnerability assessments (Croll et al. 2022; Green et al. 2025).

We ultimately selected species distribution (area of occupancy, AOO), number of occurrence records, lifespan, and the ability to reproduce with parthenogenesis, as the sensitivity variables for this assessment. Lifespan and reproductive strategies guide the recoverability of a species from anthropogenic pressures. Variables considered but not selected due to data availability included: habitat specificity, reproductive output (i.e., fecundity), body size, evolutionary distinctiveness, and dispersal ability. Where additional data is available for future biological surveys, we suggest these variables be considered for future vulnerability assessments.

The Exposure score, being a specific anthropogenic pressure on a species, was quantified as the number of times a given species was previously reported as advertised in the Australian pet trade (Lassaline et al., 2026). We assumed that a higher frequency of advertisements for a species indicates a higher demand in trade, and more individuals traded. Other variables considered for the Exposure score, but ultimately not selected due to data availability, included species value in trade (i.e., price) and the number of individuals advertised.

### 2) Assessment preparation

To identify which species in the Australian domestic invertebrate trade were short range endemics (SRE’s), we calculated their area of occupancy and extent of occurrence (EOO). Occurrence records for native Australian invertebrate species were obtained from the Global Biodiversity Information Facility (GBIF) using the R package *rgbif* (Chamberlain et al. 2017; GBIF 2026). Only records with geographic coordinates were retained, and where necessary, occurrences were spatially constrained to Australia. We also removed records with a reported coordinate uncertainty greater than 5 km. Cleaned occurrence records were then visually inspected using spatial plots to identify remaining spatial artefacts or anomalies (e.g., reported occurrences in the ocean). For each species, cleaned occurrence coordinates were used to calculate extent of occurrence (EOO) and area of occupancy (AOO) following the International Union for Conservation of Nature’s (IUCN) Red List guidelines; as implemented in the *red* package using the recommended 2 km grid size. Extent of occurrence (EOO) was calculated as the minimum convex polygon encompassing all occurrence points, while AOO was estimated using a standard grid-based approach (Cardoso 2017). For the scope of this study, the final list of short-range traded species was selected based on the threshold criteria of AOO ≤ 500 km² or EOO ≤ 5,000 km². Following IUCN’s Redlist guidelines, if EOO was less than AOO, EOO was changed to be equal to AOO to ensure consistency with the definition of AOO as an area within EOO. Ultimately, 266 out of the 586 species in trade had an AOO ≤ 500 km², classifying them as extreme short-range endemics and qualifying them for inclusion in the vulnerability framework.

Variables relating to distribution, abundance, longevity and trade were categorised on a five-level scale (1–5). Information on longevity and parthenogenesis were collected from a wide range of scientific and grey literature (Appendix 1). Information on longevity was often unavailable at the species level and was therefore estimated using genus- or family-level information where necessary. Where no longevity data were available, a score of 3 was assigned as a conservative intermediate value. These cases are indicated in the dataset (n = 47 species, 17.6%). No genus- or family-level averages or other proxy values were used for the remaining variables.

Parthenogenesis was treated as a binary variable and was categorised as either exhibiting this reproductive strategy (Rating = 0) or not (Rating = 5), to maintain consistency with the rating scale for other variables. This approach reflected the assumption that parthenogenesis either contributed to vulnerability under the framework or did not, rather than representing a scaled degree of vulnerability. Abundance, lifespan, and trade threat were manually categorised according to relative vulnerability, with category thresholds determined through researcher judgement informed by biological knowledge and the distribution of available data. Manual categorisation was used because these variables exhibited uneven and highly skewed distributions, for which equal width classification would have resulted in poorly differentiated categories.

### 3) Scoring

The relative weighting of variables constituting the Sensitivity index was determined using the analytic hierarchy process (AHP); a multi-criteria decision-making approach that derives variable weights through structured pairwise comparisons (Vaidya and Kumar 2006). The AHP was selected because its hierarchical structure aligns with the organisation of sustainability and vulnerability frameworks, and it incorporates a consistency verification check to evaluate the reliability of expert judgments (Anselin et al. 1989; Arsenault et al. 2025; Gan et al. 2017).

To develop a framework for determining the relative importance of these variables, we sought preliminary expert input from researchers and practitioners within our research network with expertise in ecology and wildlife conservation, specifically individuals with backgrounds in wildlife trade and entomology (n = 18). Contributors were asked to perform pairwise comparisons between variables using Saaty’s nine-point scale, where a score of 1 represented equal importance and a score of 9 indicated extreme importance of one variable over another (Saaty 1987). The mean importance for each variable was then calculated using a geometric mean to preserve the multiplicative relationships of each response.

Pairwise comparison matrices were constructed, and variable weights were calculated from the principal eigenvalue of the matrices. The consistency of responses was assessed by calculating the consistency ratio (CR) (Saaty 1987). A consistency ratio below 0.10 was considered indicative of acceptable consistency.

### 4) Analysis

For each species, vulnerability was calculated as the product of the Sensitivity index and the Exposure score: *Vulnerability = Sensitivity × Exposure*.

Sensitivity score was derived from four ecological variables structured within the analytic hierarchy process framework (distribution, number of recorded occurrences, longevity, and ability to reproduce through parthenogenesis), and was normalised so that the combined weight of the sensitivity component was equal to 1 (Figure 3).

**Figure 3:**
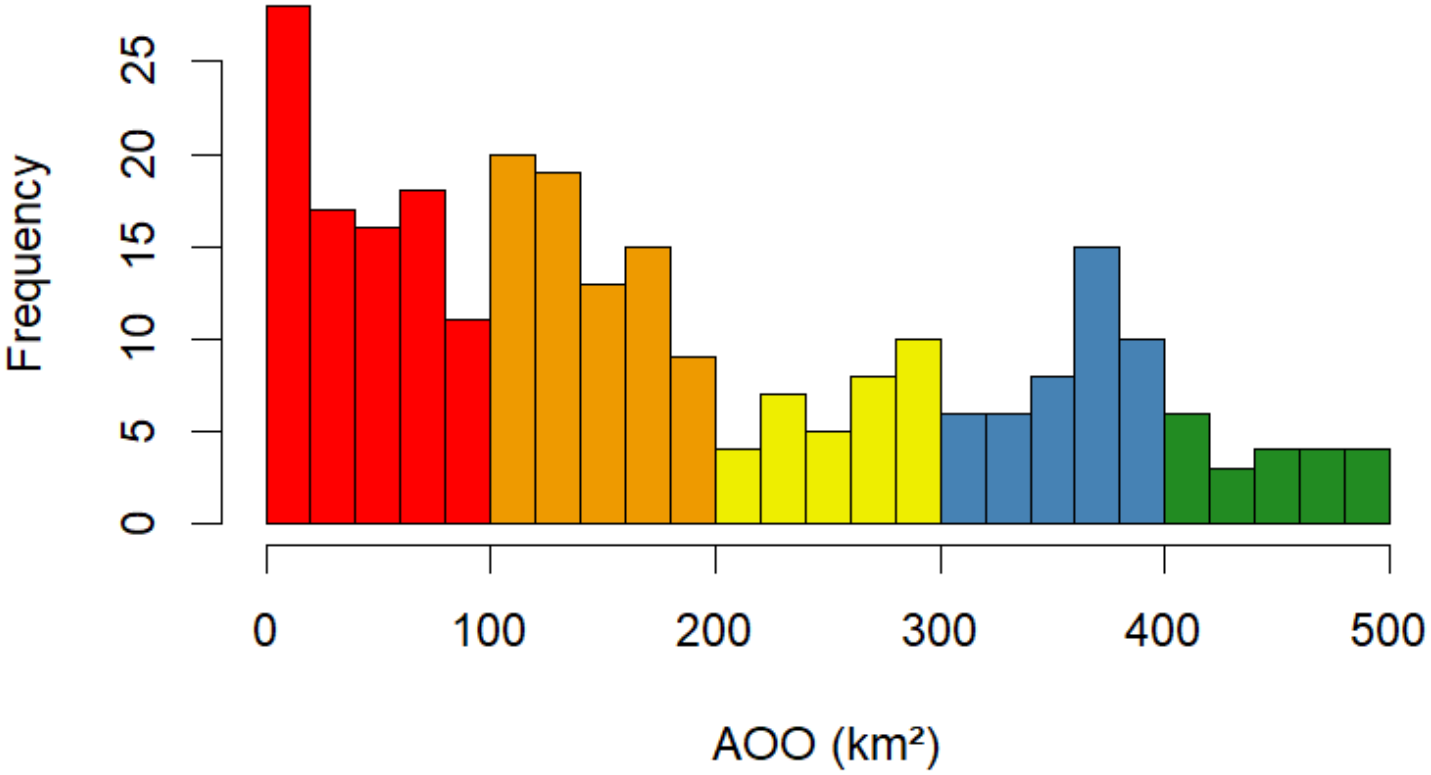
Frequency distribution of area of occupancy (AOO) for the 266 Australian terrestrial SRE invertebrate species. Colours indicate the five equal-width categories used to assign AOO ratings, ranging from 5 (red, highest vulnerability, smallest AOO) to 1 (green, lowest vulnerability, largest AOO).

All further statistical analyses were conducted in the R software for statistical and graphical computing (version 4.3.1) (R Core Team 2021). Summary statistics and plots were created using packages *dplyr* and *ggplot2* (Wickham 2016; Wickham et al. 2021). Spatial analyses were conducted using the packages *sf* and *red* (Cardoso 2017; Pebesma and Bivand 2023). To visualise broadscale spatial patterns in the distribution of SRE species, a kernel density estimate was generated from occurrence records using the packages *ggplot2* and *sf* (Pebesma and Bivand 2023). A bandwidth of 200 km was applied to both spatial dimensions, producing a smoothed representation of areas with relatively high concentrations of SRE occurrence records. A 200 km bandwidth was selected to provide a representation of spatial concentration while reducing the potential for individual sensitive occurrence localities to be inferred from the observed map. To determine whether highly vulnerable species were already afforded legislative protection, the top ranked species were cross-referenced against the list of nationally protected species under Australia’s Environment Protection and Biodiversity Conservation Act 1999 (EPBC Act). Species were also cross-referenced against relevant threatened species legislation in each Australian state and territory, and IUCN’s Red List of Threatened Species.

## Results

A large proportion of the species identified within the Australian pet trade (33.7%, n = 90) had an area of occupancy (AOO) of less than 100 km² (Figure 3). A further 28.5% of species had an AOO between 100 and 200 km², indicating that more than 60% of species assessed in this framework occupied an area of less than 200 km².

The analytic hierarchy process revealed a high level of consistency among the 18 experts, with an overall consistency ratio of 0.0076 for the pairwise comparison matrices. This value is well below the recommended threshold of 0.10, indicating that the resulting variable weightings were sufficiently consistent for subsequent analysis.

Distribution received the highest weighting score from the analytic hierarchy process (0.3686), followed by the number of occurrences, longevity and parthenogenesis (Figure 4, Table 1).

**Figure 4:**
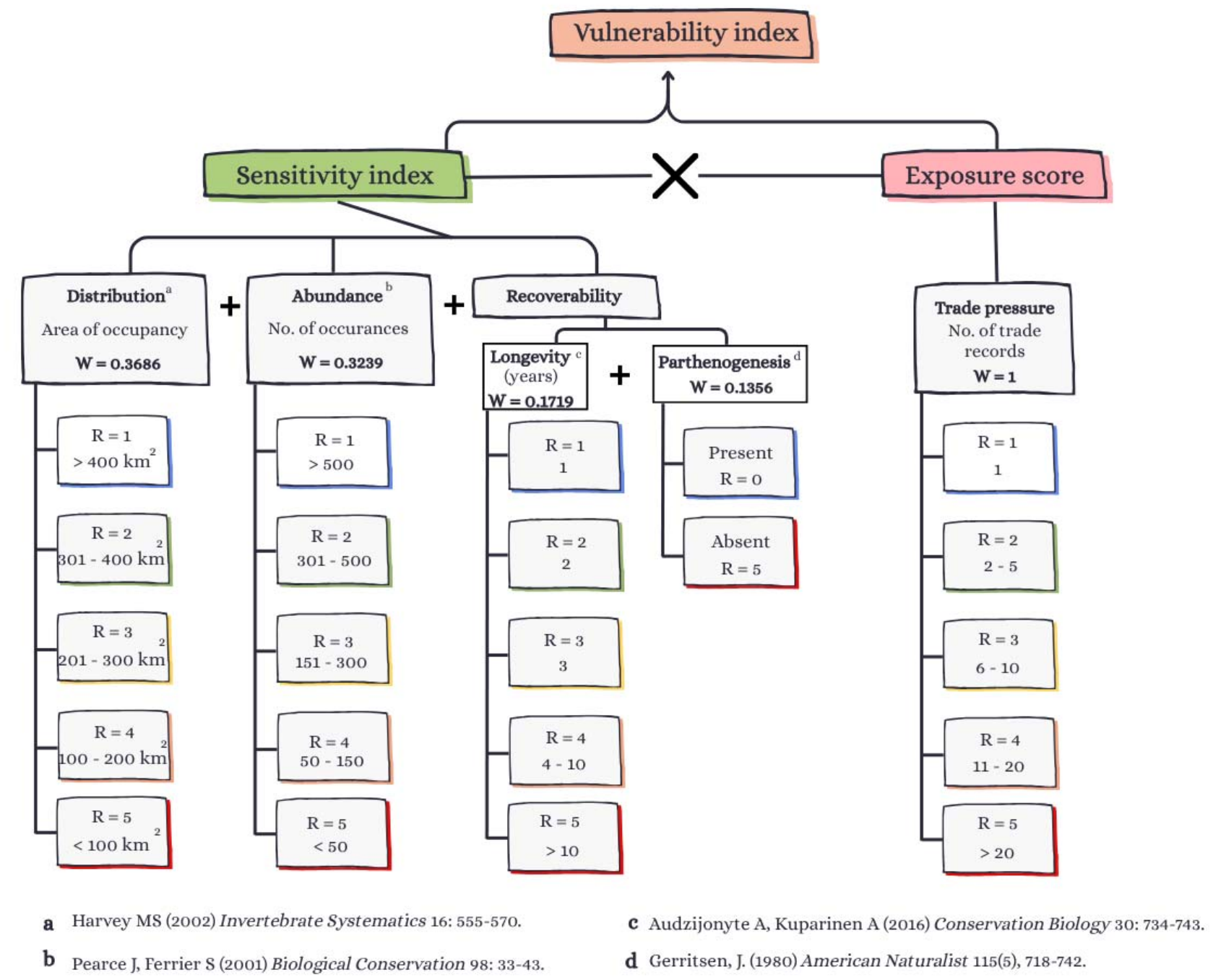
Components of the vulnerability assessment framework, including variables incorporated in the Sensitivity index, and how each variable was categorised and weighted. R refers to the rating of each category (i.e., the numeric value allocated to it), and W refers to the weight of each variable. Letters (a-d) correspond to knowledge sources that justify the inclusion of each variable. Distribution was informed by (Harvey 2002), abundance by (Pearce and Ferrier 2001), longevity by (Audzijonyte and Kuparinen 2016), and parthenogenesis by (Gerritsen 1980).

**Figure 5:**
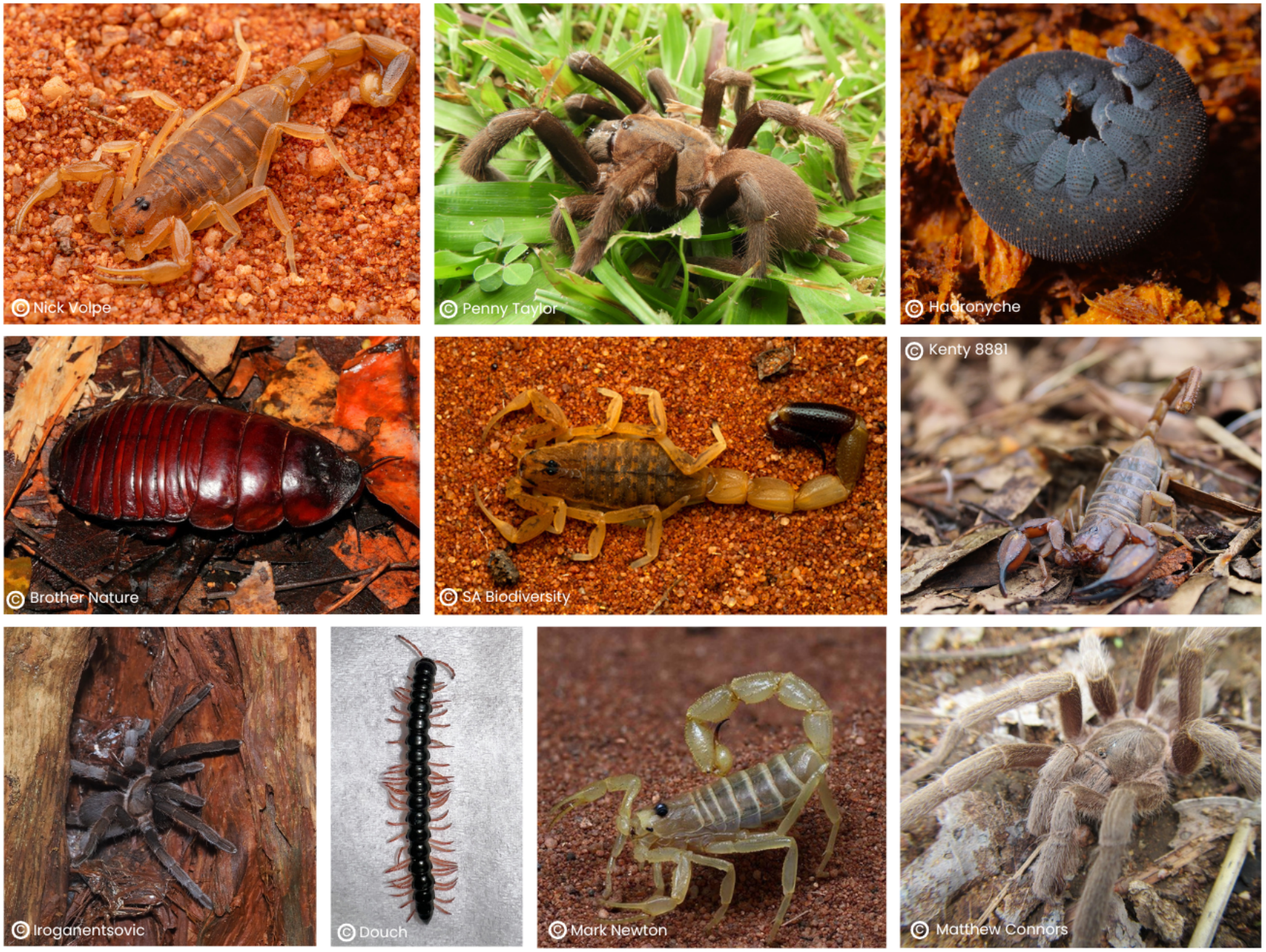
The ten highest ranking vulnerable invertebrate species. From left-to-right, top: *Lychas buchari*, *Selenocosmia crassipes, Euperipatoides rowelli*; middle: *Macropanesthia rhinoceros*, *Isometroides vescus*, *Urodacus elongatus*; bottom: *Coremiocnemis tropix*, *Heterocladosoma bifalcatum*, *Australobuthus xerolimniorum*, *Selenotypus plumipes*. Images were sourced from iNaturalist (https://www.inaturalist.org/), are research-grade identifications, and are licensed under CC BY-NC 4.0 (https://creativecommons.org/licenses/by-nc/4.0/) with attribution provided on each image.

**Table 1:** Variables included in the species vulnerability assessment framework, their ecological interpretation (units of measurement), effect on vulnerability, categorisation approach, and analytic hierarchy process derived weights. The framework is structured into two components: a Sensitivity index and an Exposure score. Categories were defined where higher values represent greater vulnerability for variables that increase risk (e.g. shorter lifespan, lower abundance), and lower vulnerability for variables that reduce risk (e.g. larger distribution). Parthenogenesis was treated as a binary variable and categorised to reflect reduced vulnerability where present.

| **Variable** | **Units** | **Effect on vulnerability** | **Mean ± SD** | **Range** | **Categories** | **Weight** |
|---|---|---|---|---|---|---|
| **Sensitivity index** |  |  |  |  |  |  |
| Distribution | Km<sup>2</sup> | Decrease | 6.169 ±1.519 | 4-496 | 5 categories<br>5 = high<br>1 = low | 0.3686 |
| No. of occurrences | No. of occurrences | Decrease | 114.69 ± 117.62 | 1-956 | 5 categories<br>5 = high<br>1 = low | 0.3239 |
| Longevity | Years | Increase | 6.179 ± 6.555 | 1-20 | 5 categories<br>5 = high<br>1 = low | 0.1719 |
| Parthenogenesis | Binary | Decrease | 4.812 ± 0.953 | - | Present = 0, absent = 5 | 0.1356 |
| **Exposure score** |  |  |  |  |  |  |
| Trade pressure | No. of advertisements | Increase | 9.677 ± 38.169 | 1-506 | 5 categories<br>5 = high<br>1 = low | 1.000 |

The number of recorded occurrences for each species ranged from single reported occurrences (e.g., *Pareremus tigrinus*) to 965 occurrences (*Urodacus armatus*). The distribution (AOO) of the SRE species ranged from 4km^2^ (e.g., *P. tigrinus*) to 496 km^2^ (e.g., *Bergama cordata*). Trade pressure, represented as the Exposure score, ranged from 1–506 advertisements (Table 1).

Ultimately, the framework identified variation in vulnerability among the 266 Australian terrestrial invertebrate species assessed, with Vulnerability indices ranging between 1.7 and 25 (Appendix 1).

Of the 266 assessed species, the 15 highest ranking species had a Vulnerability index between 18 and 25 (Table 2). Eleven of these 15 species (73.3%) were arachnids (class Arachnida), with the remaining species comprising two insects (*Macropanesthia rhinoceros, Eurycnema goliath*), one velvet worm (*Euperipatoides rowelli*), and one millipede (*Heterocladosoma bifalcatum*).

**Table 2:** The top 15 ranked vulnerable species, and their Sensitivity and Exposure scores, and calculated Vulnerability index.

| **Species name** | **Common name** | **Sensitivity score** | **Exposure score** | **Vulnerability index** |
|---|---|---|---|---|
| *Lychas buchari* | Yellow sand scorpion | 5 | 5 | 25 |
| *Selenocosmia crassipes* | Queensland whistling tarantula | 5 | 5 | 25 |
| *Euperipatoides rowelli* | Velvet worm | 4.5042 | 5 | 22.521 |
| *Coremiocnemis tropix* | Pygmy rainforest tarantula | 4.3075 | 5 | 21.5375 |
| *Isometroides vescus* | Spider-hunting scorpion | 4.3075 | 5 | 21.5375 |
| *Macropanesthia rhinoceros* | Giant burrowing cockroach | 4.3075 | 5 | 21.5375 |
| *Urodacus elongatus* | Flinders Ranges scorpion | 4.3075 | 5 | 21.5375 |
| *Heterocladosoma bifalcatum* | Red-legged millipede | 4.2876 | 5 | 21.438 |
| *Australobuthus xerolimniorum* | Salt lake scorpion | 5 | 4 | 20 |
| *Selenotypus plumipes* | Australian feather-leg tarantula | 5 | 4 | 20 |
| *Hadronyche cerberea* | Southern tree-dwelling funnel-web spider | 3.9836 | 5 | 19.918 |
| *Cercophonius kershawi* | Mallee wood scorpion | 3.9389 | 5 | 19.6945 |
| *Hadronyche levittgreggae* | Funnel-web spider | 4.6761 | 4 | 18.7044 |
| *Hadronyche versuta* | Blue mountains funnel-web spider | 3.6597 | 5 | 18.2985 |
| *Eurycnema goliath* | Goliath stick insect | 3.6344 | 5 | 18.172 |

None of the top 15 most vulnerable species (Table 2) are listed under National (EPBC Act) or State protected/threatened species lists, nor have had their conservation status assessed by the International Union for Conservation of Nature (IUCN). In the state of Queensland, permits are required for the collection and keeping of tarantula species (Family Theraphosidae), and scorpions of the *Urodacus* genus.

Occurrence records for the 266 SRE species were spatially concentrated in several regions across Australia (Figure 6). The highest relative occurrence densities were observed along the east coast of northern Queensland, and around the cities of Brisbane, Queensland, and Sydney, New South Wales. Smaller, yet still observable, concentrations of SRE records were displayed in South Australia, particularly in the vicinity of Adelaide and the northern Eyre Peninsula, and in southwestern Western Australia, around Perth. These patterns indicate that records of traded SRE species are geographically concentrated rather than uniformly distributed across the continent. With the exception of the pronounced concentration in Northern Queensland, higher relative occurrence densities were generally observed in areas surrounding capital cities. This may reflect differences in sampling effort or observer bias.

**Figure 6:**
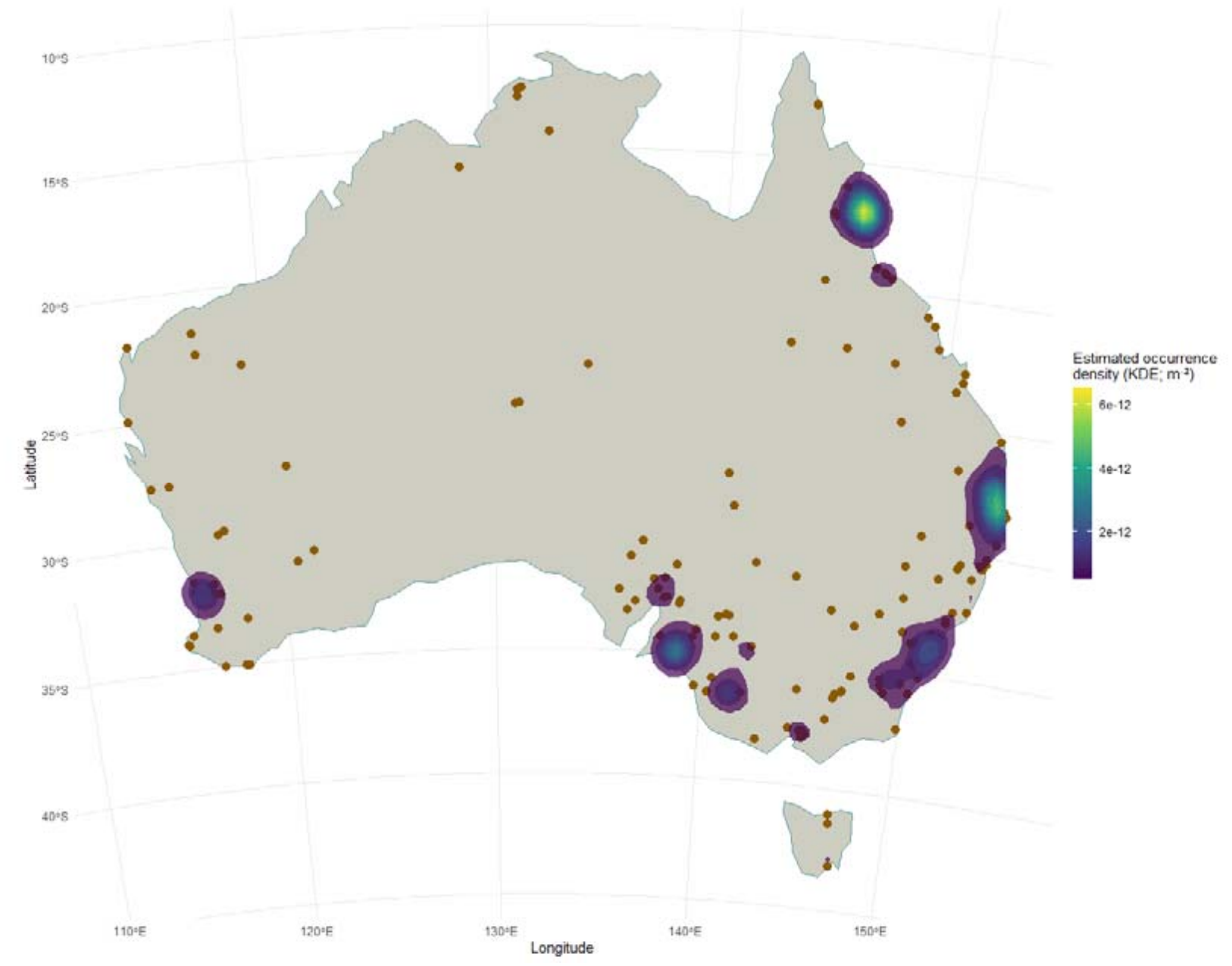
Broad-scale spatial concentration of the traded short-range endemic (SRE) species occurrence records across Australia. Points represent one randomly selected coordinate per species, while colours represent relative occurrence-record density derived using a two-dimensional kernel density estimate (m^-2^) with a 200km bandwidth. Individual occurrence localities are not shown to minimise displaying the locations of vulnerable species.

## Discussion

The increasing invertebrate pet trade is an under-recognised biosecurity and conservation threat globally. Many of the species traded are poorly recorded and are potentially at risk from unsustainable and unregulated harvest. Nearly half of the 568 species currently documented in Australia’s pet trade (47%) exhibit extreme short range endemism characteristics, with an area of occupancy less than 500 km^2^. These species are inherently more vulnerable to environmental disturbances, including wild harvest, due to their life-history traits and distribution (Hernández-Yáñez et al. 2022; McKinney 1997). We have successfully demonstrated a transparent and scalable framework for quantifying relative vulnerability among species and triaging conservation action, which is capable of rapidly prioritising species for further assessment.

This assessment methodology is particularly important for terrestrial invertebrates, where comprehensive species-level threat assessments are often impractical given their vast diversity and the large number of under-studied taxa (Cardoso et al. 2011). Globally, invertebrate declines and extinctions often remain undetected, with ‘ghost extinctions’ arising from the disproportionate number of undescribed taxa and comparatively limited research attention directed towards them (Woinarski et al. 2024). There is therefore a need for rapid and adaptable approaches that can identify potentially vulnerable species before population declines become irreversible. This study represents the first attempt to systematically integrate wildlife trade exposure and biological sensitivity into a screening framework for Australian terrestrial invertebrates. Although intended as a preliminary assessment, the framework identified several species and taxonomic groups that warrant further investigation and monitoring.

### Most vulnerable Australian taxa

Our vulnerability rankings demonstrated that trade demand and vulnerability are not always synonymous. Frequently traded species are not necessarily the most vulnerable to exploitation, and species with relatively lower levels of recorded trade may still warrant attention where their biological sensitivity is high. This reflects the main concept behind this framework: vulnerability can emerge from the interaction between exposure to wildlife trade and biological characteristics that determine a species capacity to withstand and recover from exploitation. Consequently, conservation prioritisation based solely on trade frequency could overlook species for which relatively low levels of wild collection may have substantial ecological consequences.

Arachnids were disproportionately represented among the species with the highest vulnerability scores in this assessment. This does not suggest that all arachnids are inherently more vulnerable than other invertebrate groups, but instead reflects the interaction between biological, geographic and anthropogenic variables that may increase their susceptibility to over-exploitation. Short-range endemism is particularly prevalent among Australian arachnids, including scorpions and mygalomorph spiders, which are often characterised by restricted distributions, limited dispersal ability, and slow life-history traits. These characteristics can increase vulnerability to wild harvesting because populations have limited capacity to recover following localised depletion. At the same time, trade pressure may be concentrated on distinct arachnid groups, such as scorpions and tarantulas, because of their popularity among collectors (Lassaline et al. 2026b; Marshall et al. 2022). Species perceived as dangerous and unique often hold particular appeal in the invertebrate trade, potentially increasing demand for arachnids relative to other invertebrate group (La Laina et al. 2021; Lassaline et al. 2023). The concentration of highly vulnerable arachnid species may reflect the convergence of high trade exposure with biological characteristics that constrain population recovery. Where the success of captive breeding of particular species is limited by specialised behavioural or environmental requirements, continued demand for wild sourced individuals is even more pronounced.

We recommend that Australian arachnids receive future monitoring and detailed assessments of conservation impacts. None of the species identified as having the highest vulnerability scores are afforded protection under Australian national or state threatened species legislation, potentially highlighting gaps in current conservation priorities. Further applications of the framework, incorporating more comprehensive biological variables and a larger pool of experts for the analytic hierarchy process, would help evaluate the robustness of these patterns.

### Future framework applications

Our vulnerability assessment framework is designed to be transferrable across other taxonomic groups affected by wildlife trade. As a screening and prioritisation framework, its purpose is not to replace detailed species assessments, but to provide a rapid and transparent means of identifying species that warrant further investigation and protection most urgently. The frameworks application requires three broad categories of information: (i) trade pressure, (ii) distribution and abundance, and (iii) life-history traits. First, data on the frequency of trade advertisements are required to quantify trade pressure (i.e., the Exposure score) for each species. Within Australia alone, numerous existing wildlife trade datasets could be used to apply the framework to other taxonomic groups (Chekunov et al. 2026; Hill et al. 2024; Maher et al. 2023; Toomes et al. 2023). The availability of trade datasets varies considerably between regions globally and are largely located within either peer-reviewed literature or government funded surveillance datasets. Second, occurrence data are required to estimate species’ distributions and relative abundance. Such data are readily available through public biodiversity databases, including the Atlas of Living Australia (ALA) and the Global Biodiversity Information Facility (GBIF). Similarly, the availability and completeness of biodiversity occurrence and life-history data vary between regions. Third, information on species life-history data is required. As we have demonstrated, these data can be obtained from a range of sources, including scientific literature, biodiversity repositories, and wildlife trade platforms and forums.

Of the three required data categories, life-history information may present the greatest challenge to acquire, particularly for poorly studied taxa. However, with the combination of other variables, the framework allows flexibility in the selection of life-history traits and, where species-specific information is unavailable, informed generalisations can be made. For example, when longevity data could not be obtained for several species in our worked example, species were assigned to the intermediate category for this variable (i.e., a score of 3). This flexibility facilitates the application of the framework across taxonomically diverse groups, while importantly still allowing preliminary prioritisation to be undertaken for data deficient species. Furthermore, the framework is not dependant on a fixed set of life-history variables, allowing for included traits to suit the biological characteristics of the assessed group. For example, parthenogenesis would not be relevant to an assessment of bird species, but could be replaced by alternative traits, such as reproductive output or dispersal ability.

Social influences could also be incorporated into future assessments, as societal behaviour and popular culture can influence wildlife trade demand and consequently shape trade dynamics. These influences are rarely documented systematically and are not commonly incorporated into conservation prioritisation frameworks, despite their ability to rapidly change consumer interest and subsequent trade pressure on particular taxa. Popular media can increase public awareness of wildlife trade and increase interest in particular species, with the 2003 Disney film *Finding Nemo* often involved in discussions surrounding the influence of popular culture on demand for aquarium species (Militz and Foale 2017). Incorporating indicators of public interest, such as Google search trends, could therefore provide a measure of potential social drivers of trade demand in future assessments (Valdez 2021).

By communicating relative vulnerability to trade, the outcomes from our framework could help wildlife keepers make more informed and ethical decisions about the species they acquire. This could encourage more responsible purchasing practices and support broader awareness of the conservation implications of invertebrate trade.

### Implications for biodiversity data management

Additionally, vulnerability could be used to inform broader biodiversity data sharing decision making. We recommend that calculated Vulnerability indices be used to inform decisions regarding which species should be included on sensitive species lists. Such approaches could complement, rather than replace, existing threatened-species assessments by directing limited monitoring and research resources towards species where the consequences of continued exploitation may be greatest.

Species identified as highly vulnerable to overexploitation through wildlife trade may require greater protection of their occurrence data, including the generalisation or omission of precise locality information from public databases and publications. This would reduce the risk that publicly accessible occurrence data could be used to identify populations or collection sites and facilitate targeted wild collection. These measures should be applied strategically, balancing the need to protect vulnerable species from exploitation with the importance of maintaining access to biodiversity data for legitimate research and conservation purposes.

### Future recommendations

As this trait-based vulnerability framework does not require extensive modelling capabilities or statistical expertise, it can be applied quite rapidly to large taxonomic groups. However, there are limitations to this framework, both with the methods used to assess vulnerability, and the data used to populate it. Importantly, the framework assumes that the interaction between biological sensitivity (i.e., a species ability to recover from over-exploitation) and trade exposure provides a proxy for vulnerability to exploitation, but the extent to which these impacts translate into conservation impacts remains uncertain. Here, we provide six recommendations to address these limitations.

#### Recommendation 1: Addition of wildlife trade volume data

The trade data captured only represented observable wildlife trade, meaning that undetected, covert aspects of the Australian invertebrate trade are likely excluded and trade pressure may be highly underestimated. Trade advertisements also provide a measure of market exposure rather than the number of individuals actually traded, and therefore cannot directly quantify harvested volumes. Where available, wildlife trade volume data would also provide a useful addition to future vulnerability frameworks.

The trade data utilised for this assessment were collected through dedicated research efforts, rather than an established government-led system for monitoring wildlife trade. In Australia, as in many other countries, there is a strong reliance on independent research to generate information that is otherwise largely unavailable to practitioners and regulators. We recommend the continued investment in research-led wildlife trade surveillance to generate the data needed to support frameworks such as ours.

#### Recommendation 2: Incorporating sourcing practices

The framework does not explicitly distinguish between traded individuals that are captive bred or sourced from wild populations. While trade frequency can reflect species demand, it does not directly reflect wild harvesting pressure, and some species may be captive bred.

Consequently, all species with high trade demand may not necessarily experience the same harvesting pressures. Additionally, where captive breeding is limited and trade is known to be predominantly supplied by wild-collected individuals, trade exposure may represent a substantially greater conservation risk. The absence of information on the source of species therefore introduces uncertainty into the Exposure score. Future applications of the framework should incorporate information on sourcing practices where possible.

#### Recommendation 3: Incorporating data confidence

Biological data may also introduce uncertainty into the framework. Species identifications, distribution records and life-history traits such as longevity may be incomplete or uncertain, particularly for poorly studied taxa. Although the framework allows informed generalisations where species-specific information is unavailable, assigning intermediate scores in the absence of data may reduce the ability to distinguish genuinely intermediate vulnerability from uncertainty. Future applications could incorporate explicit measures of data confidence or uncertainty alongside vulnerability scores.

#### Recommendation 4: Expanding and diversifying the analytic hierarchy process expert panel

While the analytic hierarchy process provides a transparent and repeatable mechanism for incorporating expert knowledge, it inevitably reflects the perspectives of the participating experts. Future applications should involve larger and more taxonomically diverse expert panels from different research and geographic backgrounds, sourced through a more systematic recruitment process. This may help mitigate potential biases and produce more representative weighting values.

#### Recommendation 5: Validating the Vulnerability index against available data

The Vulnerability index has not been validated against observed population trends. However, this is particularly difficult for data deficient terrestrial invertebrates. Therefore, the framework identifies relative vulnerability rather than demonstrating that the highest ranked species are currently undergoing population declines as a result of wildlife trade. Long term population studies and targeted field studies could be incorporated into future assessments as an important next step to confirm species vulnerability.

#### Recommendation 6: Accounting for interacting threats in future assessments

Wildlife trade is unlikely to represent the sole threat faced by most species identified by the framework. Rather, over-exploitation through wildlife trade may interact synergistically with existing pressures, including habitat loss, climate change, invasive species, pollution and pesticide exposure (Isbell et al. 2023). Vulnerability is therefore likely to arise from the interaction between trade exposure and biological sensitivity, rather than either factor alone. The framework should therefore be viewed as identifying species for which wildlife trade may represent a significant additional pressure, rather than as an assessment of overall extinction risk.

## Conclusion

We have demonstrated the potential for rapid, trait-based screening to identify species that may be particularly vulnerable to wildlife-trade exploitation despite limited biological and conservation information. Our framework does not provide a substitute for formal threat assessments, population monitoring or species-specific investigation, nor does a high vulnerability score demonstrate that trade is currently causing population decline. Rather, it provides a transparent and repeatable means of identifying where these more resource-intensive activities may be most urgently warranted. For a taxonomic group in which majority of species remain poorly studied, such as invertebrates, this type of prioritisation may be essential to ensuring that conservation action is not restricted to species that are simply better researched. As the first study of its kind to systematically integrate wildlife trade exposure and biological vulnerability into a screening framework for wildlife, this framework provides a practical first step to more proactive and precautionary management of wildlife trade and species conservation.

## Acknowledgements

The authors acknowledge the Kaurna people as the Traditional Owners of the land where we live, learn and work. We acknowledge the Kaurna people as the custodians of the Adelaide region, and we respect and value their past, present and ongoing connection to the environment and cultural beliefs. We thank the 18 researchers from Adelaide University who assisted with the analytic hierarchy process, and all collaborators who assisted with collecting wildlife trade data used in our framework, originally published by Lassaline et. al (2026). PC is an Australian Research Council Industry Laureate Fellow (IL230100175; ‘Combatting wildlife crime and preventing environmental harm’). We also thank the Wildlife Crime Research Hub for their ongoing support with this research.

